# Antimicrobial interactions between the phytoextracts of *Callistemon citrinus* and *Eriobotrya japonica* against *Streptococcus mutans*

**DOI:** 10.1101/2021.08.13.456290

**Authors:** J. S Kavuo, Jerald Kutosi Namboko, Samuel Bukusuba, Healy Onen

## Abstract

*Streptococcus mutans* is a gram-positive bacterium in the oral cavity that is most implicated in the dental caries progression. The condition is very expensive to manage and the most commonly used products such as fluoride tooth pastes and alcohol-based mouth washes are associated with many side effects. The current study therefore focused on providing a scientific evidence to guide the use of a combination of *Eriobotrya japonica* (EJ) and *Callistemon citrinus* (CC) as actives in development of an effective and cheaper herbal formulation for management of dental caries. The objective of this study was to determine antimicrobial interactions of *Eriobotrya japonica* and *Callistemon citrinus* phytoextracts proportions against *Streptococcus mutans* bacteria. The leaves of both plants (EJ and CC) were shade-dried and pulverized into a coarse powder which were then cold macerated using ethanol (60 %) for 24 h. Phytochemical screening was conducted for the two dry extracts obtained after fan drying before they were mixed in to five different proportions (1:0, 3:1, 1:1, 1:3 and 0:1). Minimum inhibitory concentrations (MIC) and minimum bactericidal concentration (MBC) assays against *Streptococcus mutans* were done for all the proportions above with ciprofloxacin and 2.5 % Dimethyl sulfoxide (DMSO) as the positive and negative controls respectively. Antimicrobial interactions between the two extracts were also evaluated using Fractional Inhibitory and Bacterial Concentration Indices (FICI/FBCI). Results showed that EJ and CC had percentage yields of 20.05 % and 15.45 % respectively. All the extracts showed similar phytochemical profiles. They also demonstrated an inhibitory effect on *Streptococcus mutans with* MIC and MBC values ranging from 521 to 3333 μg/ml and 1042 to 3667 μg/ml respectively. However, CC: EJ (1:0) had the lowest MIC and MBC comparable to that of the standard drug at P <0.05. The FICI/FBCI were between 1.5 and 3.917. Therefore, CC: EJ (1:0) proportion markedly demonstrated better antimicrobial activity against the test organism and there are no beneficial antimicrobial interactions between the two plant extracts to inform their combination as actives for dental caries product formulation.

## Introduction

*Streptococcus mutans* is a Gram-positive bacterium in the oral cavity. It is considered as the most relevant bacteria in the progression of dental caries [1]. *Streptococcus mutans* has multiple mechanisms to colonize the tooth surface and form bacterial plaque biofilm [2]. The biofilm plaque serves as a physical barrier which limits penetration of antimicrobial agents into the deep layers of biofilm [3]. The bacteria also produce organic acids through various carbohydrate metabolism processes that dissolve tooth enamel and dentine over time leading to the development of dental caries [4–5].

Dental caries is one of the most prevalent and consequential oral diseases globally [6]. In Uganda, the overall caries prevalence was highest in adults than in children in randomly selected urban areas [7]. In advanced states, dental caries involves the pulp of the tooth and destroy tooth structure leaving only root fragments that can lead to ulcerations and abscesses [8].

According to Young & Featherstone [9], some of the methods of controlling tooth decay include: mouth saliva which maintains the oral pH, thus preventing demineralization of teeth enamel; artificial sealants which protects tooth surface from acidic environment; antibacterial mouth rinses and Fluorides. However, when cavitated carious lesions develop, they are likely to progress and require restorative treatment as part of the caries management for that patient [10]. In addition, anti-biotics such as metronidazole, B-lactam antibiotics, tetracyclines have been used and these have led to anti-microbial resistance [11]. Also, some of the ingredients used in the formulation of oral care products such as fluorides [12] and alcohol [13] have been associated with some negative effects.

Medicinal plants are currently in considerable significance view due to their special attributes as a large source of therapeutic phytochemicals that may lead to the development of novel drugs [14]. They have been reported with several therapeutic uses including antimicrobial, antidiabetic, antiviral, anticancer and antifungal activities [15]. The ones that have activity against oral microbes and have been used in treatment of oral diseases include Bloodroot, Caraway, Chamomile, Echinacea, Myrrh, Peppermint, Rosemary, Sage, Thyme, Aloe Vera, Propolis, Alfalfa, Anises, Annatto, Black cohosh, Burdock, Elderberry, Ginseng, Goldenseal and Cloves [16].

*Eriobotrya japonica* (Loquat) and *Callistemon citrinus* (Bottlebrush) have also showed promising activity against oral microbials responsible for dental carries with zones of inhibition of 4mm and 8mm respectively [17]. According to Seong [18], *Eriobotrya japonica* showed no side effects in both male and female rats thus is considered a safe traditional medicine for clinical application. A study by Bhushan [19] also showed that *Callistemon citrinus* is well tolerated and thus safe for use.

Based on the safety profile and efficacy of *Eriobotrya japonica* and *Callistemon citrinus* already documented according to Bhushan and Seong This study therefore, aimed at exploring possible beneficial anti-microbial interactions between the two plant phytoextracts against *Streptococcus mutans* to inform future herbal product formulations from the combined plant extracts.

## Materials and methods

Some of the materials, reagents and equipment used included distilled Water, Analytical Weighing Balance, vortex mixer, Oven, Autoclave, Desitmat, Whatman filter Paper No. 1, Petri Dishes, Automated pipette, pipettes tips(1000μl and 100μl), 96-well microtiter plates, Bijor bottles, cotton swaps, Ciprofloxacin(500mg), Cock borer, Ethanol, Muller Hilton broth, Blood Agar, Muller Hilton Broth, Aluminium foil, Ferric Chloride Solution, Ammonia Solution, pH Meter, Test Tubes, Flacon tubes, Mortar and Pestle, Rotary Shaker, plastic loop, Drangendoff’s Reagent, Fehling Solution, Sudan III reagent, 10% Dimethyl sulfoxide (DMSO).

### Study design and area

The study was an in-vitro experimental study carried out at Mbarara University Pharmaceutical Chemistry Analytical Research Laboratory (MUPCARL) and Epicentre Microbiology Laboratory Mbarara, Uganda.

### Preparation of plant parts

The leaves of *Eriobotrya japonica* and *Callistemon citrinus* were collected from a selected part of Mbarara University (latitude and longitude −0.61723342, 30.65666333 for *Callistemon citrinus* and −0.61670506, 30.65656174 for *Eriobotrya japonica)*. The plant parts were identified by Dr. Olete Eunice a botanist at Faculty of Science, Mbarara University of Science and Technology and issued plant identification numbers (Healy Onen 001 and Healy Onen 002 for EJ and CC respectively). Fresh leaves of *Eriobotrya japonica* and *Callistemon citrinus* were dried in a well aerated room under shade at room temperature. The dried plant materials of both plants were pulverized to a coarse powder and stored awaiting extraction.

### Cold maceration of *Eriobotrya japonica* and *Callistemon citrinus*

The coarse powder (300 g) was placed in a 2.5 L amber bottle and extracted using 1500 ml of 60 % ethanol for 3 days with vigorous shaking after a day. After 3 days, the extract was filtered using a muslin cloth and the filtrate concentrated and dried by using a fan. The final dry extract was weighed using an analytical weighing balance and percentage yield was then calculated. The extract was refrigerated at −4 °C awaiting antimicrobial studies [20].

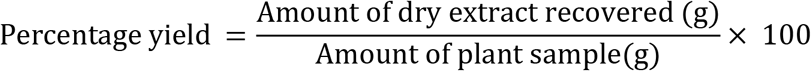

### Phytochemical Screening

The different extracts were then subjected to qualitative phytochemical tests for Alkaloids, Flavonoids, Saponins, Tannins and Essential oils [21].

### Anti-microbial Studies

#### Test organism

The Standard strain of *Streptococcus mutans* ATCC 25175 (cariogenic bacteria) was obtained from Makerere university, College of Veterinary medicine.

#### Preparation, storage of the testing reagent

Resazurin was prepared at 0.015 % by dissolving 0.015 g in 100ml of sterile water, vortexed and filter sterilized (0.22 μm filter) and stored at 4 °C for a maximum of 2 weeks after preparation [22].

#### Preparation of plant extracts and standard drug

The dry extracts from both plants were mixed in five different proportions (1:3, 1:1, 3:1, 1:0, and 0:1) by weighing 1g of the extracts separately and dissolving each in 10 ml DMSO (10 %) to produce concentration of 0.1 g/ml. 80 μl was pipetted from the above stock solution and topped up to 1ml using sterile water to produce test concentration of 80 mg/ml and this was done for each of the plant extract proportions. For Ciprofloxacin (standard drug), 400 mg of drug was dissolved in 10 ml of sterile water to obtain concentration of 40 mg/ml. 500 μl was then pipetted from this stock solution and topped up to 1ml using sterile water to form a test concentration of 0.02 mg/ml.

#### Preparation of standardized inoculum

The bacteria were sub-cultured on a blood Agar and incubated at 37 °C for 24 hours [23]. A cotton swap was used to inoculate the colonies in sterile normal saline (0.85 %) tube to obtain a McFarland standard streptococcus suspension of 0.5 at OD600, which is equivalent to 1.5×108 CFU/ml [24]. Approximately 5 ×105 CFU/ml was then prepared by carrying 1:100 broth dilution. The inoculum prepared was dispensed into the wells in less than 15 min.

#### Determination of the minimum inhibitory concentration (MIC)

50 μl of broth was dispensed in each well (1-12 and A-G) on the plate and 50 μl of extracts and standard drug was added in the first column (A-G). Serial dilutions were then carried out to obtain final concentrations in the range of 20000 μg/ml to 39.0625 μg/ml for extracts and 5 μg/ml to 0.009765625 μg/ml for the standard drug after addition of the standardized bacterial suspension (50 μl). The solvent DMSO (2.5 % v/v) was used as a negative control. The plates well then incubated in the oven at 37 °C for 24 hours30 μl of . 0.015 % resazurin was added to all wells and further incubated for 4 hours for the observation of colour change. The lowest concentration of no colour change from blue to pink (blue resazurin colour remained unchanged) was to be scored as the MIC value.

#### Determination of the minimum bactericidal concentration (MBC)

20μl of the contents from the wells before the MIC value were drawn and inoculated on a fresh blood agar incubated in an oven at 37 °C for 24 h. The minimum concentration with no colony growth was taken as the MBC [25].

#### Determination of antimicrobial interactions

Fractional inhibitory/bactericidal concentration indices (FICI/FBCI) were used as reported previously by Pratoomsoot and Gohil [26–27] to determine the presence or absence of antimicrobial interactions as shown below:

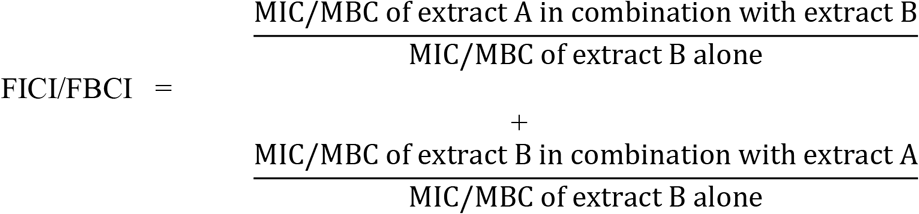

#### Data presentation and statistical analysis

The data was summarized into mean ±SED and presented in suitable tables and graphs. Statistical tools such as one-way Anova was used Turkey’s Multiple Comparison Test . The differences were statistically significant when P≤0.05. Graph-PAD Prism (version 8.0.2) software was used to conduct the data analysis.

## Results

### Percentage Yield

The percentage yield of *Eriobotrya japonica* was higher than that of *Callistemon citrinus*

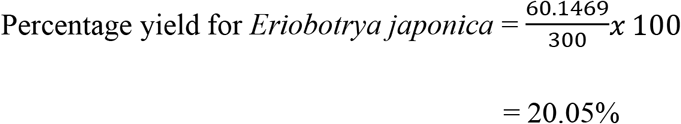

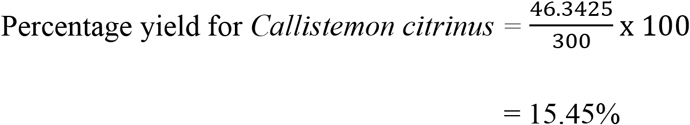

### Phytochemical Screening

The extract showed similar phytochemical groups (Table 1 and 2).

**Table 1:** showing the different phytochemicals in the ethanoic extracts of *Callistemon citrinus*

| PHYTOCHEMICAL GROUP | CONCLUSION |
| --- | --- |
| Alkaloids | - |
| Flavonoids | ++ |
| Saponins | + |
| Tannins | + |
| Volatile oils | - |

**Table 2:**
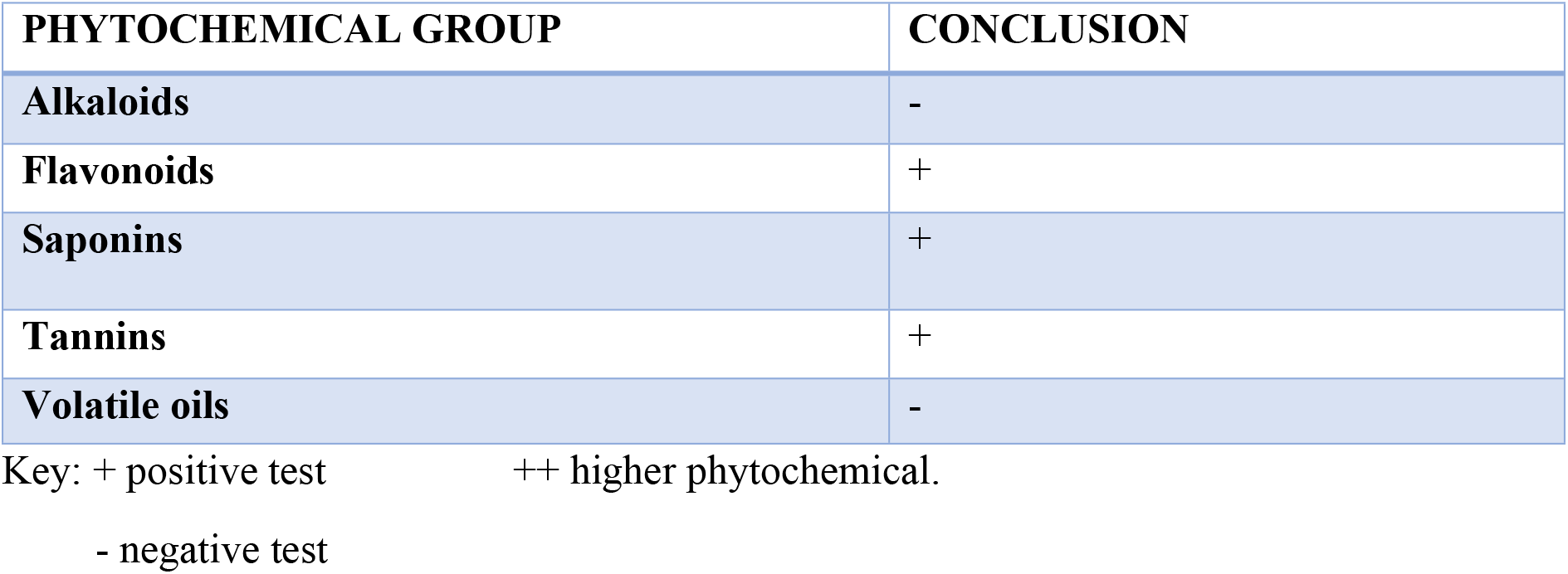
showing the different phytochemicals in the ethanoic extracts of Eriobotrya japonica

**Table 3:** showing the MIC and MBC of standard drug and plant extract.

| EXTRACT<br>AND<br>STANDAR<br>D DRUG | PLATE 1 |  | PLATE 2 |  | PLATE 3 |  | AVERAGE |  |
| --- | --- | --- | --- | --- | --- | --- | --- | --- |
|  | MIC±SED | MBC±SED | MIC±SED | MBC±SED | MIC±SED | MBC±SED | MIC±SED | MBC±SED |
| **EJ:CC (1:0)** | 2500±1443.37 | 5000±2886.75 | 2500±1443.37 | 5000±2886.75 | 5000±1443.37 | 10000±2886.75 | 3333±1443.37 | 3667±2886.75 |
| **EJ:CC (3:1)** | 1250±0 | 2500±0 | 1250±0 | 2500±0 | 1250±0 | 2500±0 | 1250±0 | 2500±0 |
| **EJ:CC (1:1)** | 625±0 | 1250±0 | 625±0 | 1250±0 | 625±0 | 1250±0 | 625±0 | 1250±0 |
| **CC: EJ (1:0)** | 312.5±180.42 | 625±360.843 | 312.5±180.42 | 625±360.843 | 625±180.42 | 1250±360.843 | 417±180.42 | 833±360.843 |
| **CC: EJ (3:1)** | 625±180.42 | 1250±360.84 | 312.5±180.42 | 625±360.84 | 625±180.42 | 1250±360.84 | 521±180.42 | 1042±360.84 |
| **Ciprofloxacin** | 0.625±0 | >1.25±0 | 0.625±0 | >1.25±0 | 0.625±0 | >1.25±0 | 0.625±0 | >1.25±0 |

**Table 4:** showing the mean difference SED, significancy and adjusted p-value of MICs and MBCs of standard drug and plant extract.

| Turkey Multiple Comparison Test (TMCT) | Mean Diff. ±SED(MIC) | Mean Diff. ±SED(MBC) | Significance (MIC) | Significance (MBC) | P-value (MIC) | P-value (MBC) |
| --- | --- | --- | --- | --- | --- | --- |
| **Cp Vs EJ:CC (1:0)** | -3332±85.05 | -6665±977.2 | *** | *** | 0.0002 | 0.0002 |
| **Cp Vs EJ:CC (3:1)** | -1249±85.05 | -2499±977.2 | No | No | 0.1825 | 0.1821 |
| **Cp Vs EJ:CC (1:1)** | -623.8±85.05 | -1249±977.2 | No | No | 0.7918 | 0.7911 |
| **Cp Vs CC: EJ (3:1)** | -519.6±85.05 | -1040±977.2 | No | No | 0.8866 | 0.8861 |
| **Cp Vs CC: EJ (1:0)** | -415.4±85.05 | -832.1±977.2 | No | No | 0.9515 | 0.9512 |
| **EJ:CC (1:0) Vs EJ:CC (3:1)** | 2083±85.05 | 4167±977.2 | * | * | 0.0109 | 0.0109 |
| **EJ:CC (1:0) Vs EJ:CC (1:1)** | 2708±85.05 | 5417±977.2 | ** | ** | 0.0014 | 0.0014 |
| **EJ:CC (1:0) Vs CC: EJ (3:1)** | 2813±85.05 | 5625±977.2 | *** | *** | 0.0010 | 0.0010 |
| **EJ:CC (1:0) Vs CC: EJ (1:0)** | 2917±85.05 | 5833±977.2 | *** | *** | 0.0007 | 0.0007 |
| **EJ:CC (3:1) Vs EJ:CC (1:1)** | 625.0±85.05 | 1250±977.2 | No | No | 0.7905 | 0.7905 |
| **EJ:CC (3:1) Vs CC: EJ (3:1)** | 729.2±85.05 | 1458±977.2 | No | No | 0.6751 | 0.6751 |
| **EJ:CC (3:1) Vs CC: EJ (1:0)** | 833.3±85.05 | 1667±977.2 | No | No | 0.5530 | 0.5530 |
| **EJ:CC (1:1) Vs CC: EJ (3:1)** | 104.2±85.05 | 208.3±977.2 | No | No | >0.9999 | >0.9999 |
| **EJ:CC (1:1) Vs CC: EJ (1:0)** | 208.3±85.05 | 416.7±977.2 | No | No | 0.9977 | 0.9977 |
| **CC: EJ (3:1) Vs CC: EJ (1:0)** | 104.2±85.05 | 208.3±977.2 | No | No | >0.9999 | >0.9999 |

**Table 5:** showing the FICI/FBCI of the different extract proportions

| Extract proportions | FICI(Mean ±SED) | FBCI(Mean ±SED) |
| --- | --- | --- |
| EJ:CC (3:1) | 3.833±1.155 | 3.917±1.465 |
| EJ:CC (1:1) | 1.875±0.649 | 1.875±0.649 |
| CC: EJ (3:1) | 1.5±0.649 | 1.5±0.649 |

### MIC And MBC

Ciprofloxacin shows the lowest MIC and MBC while EJ:CC (1:0) having the highest MIC/MBC values (Fig 2 and 3 respectively). The P-value (0.0002) obtained when comparing Ciprofloxacin and *Eriobotrya japonica* was significant whereas the P-value (0.09155) obtained when comparing Ciprofloxacin and *Callistemon citrinus* was not significant. Increasing the quantity of *Eriobotrya japonica* in the proportions increased the MIC/MBC values whereas increasing the quantity of *Callistemon citrinus* in the proportions increased, decreases the MIC/MBC values.

**Figure 1:**
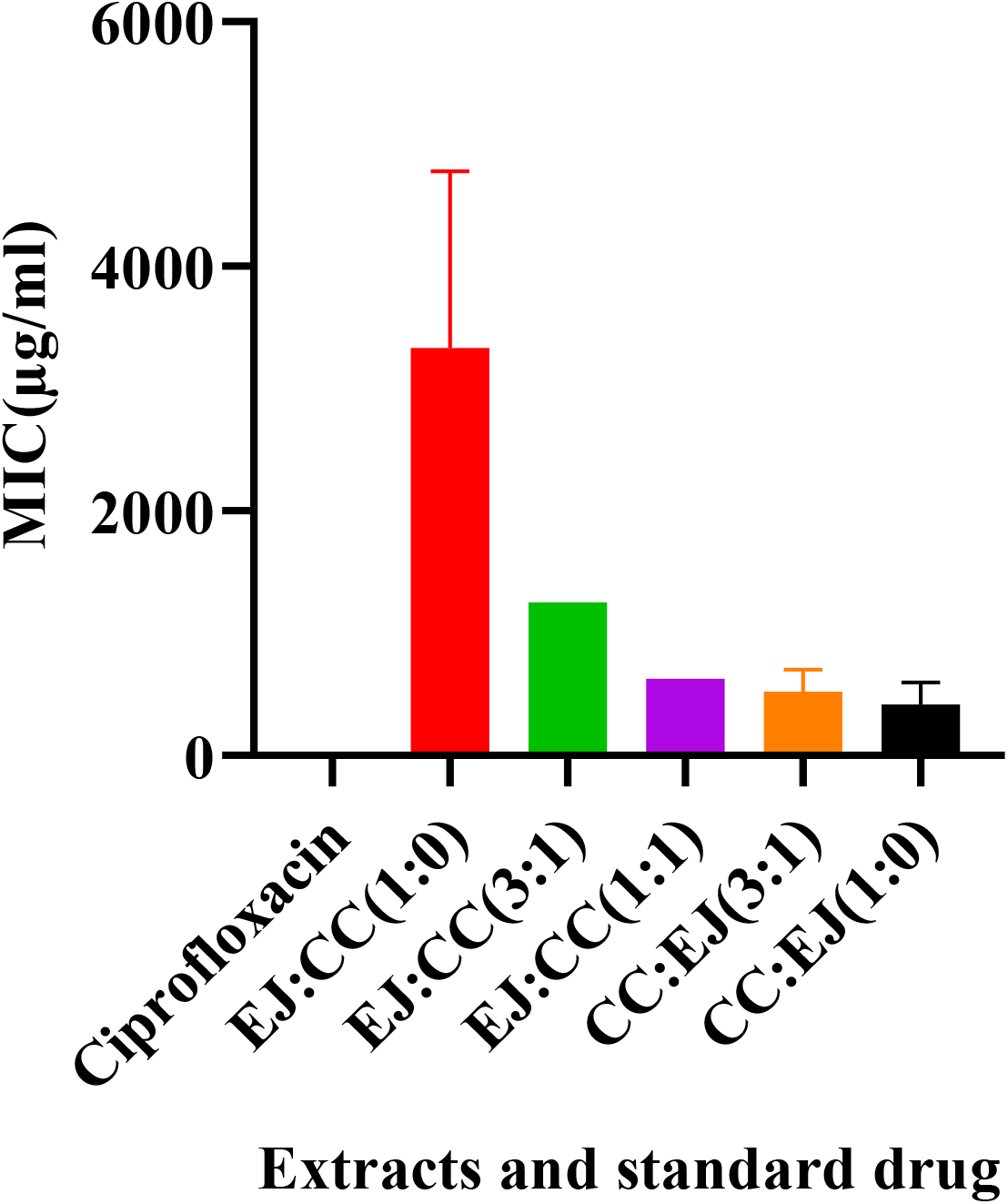
showing the MICs of different extract proportions and standard drug.

**Figure 2:**
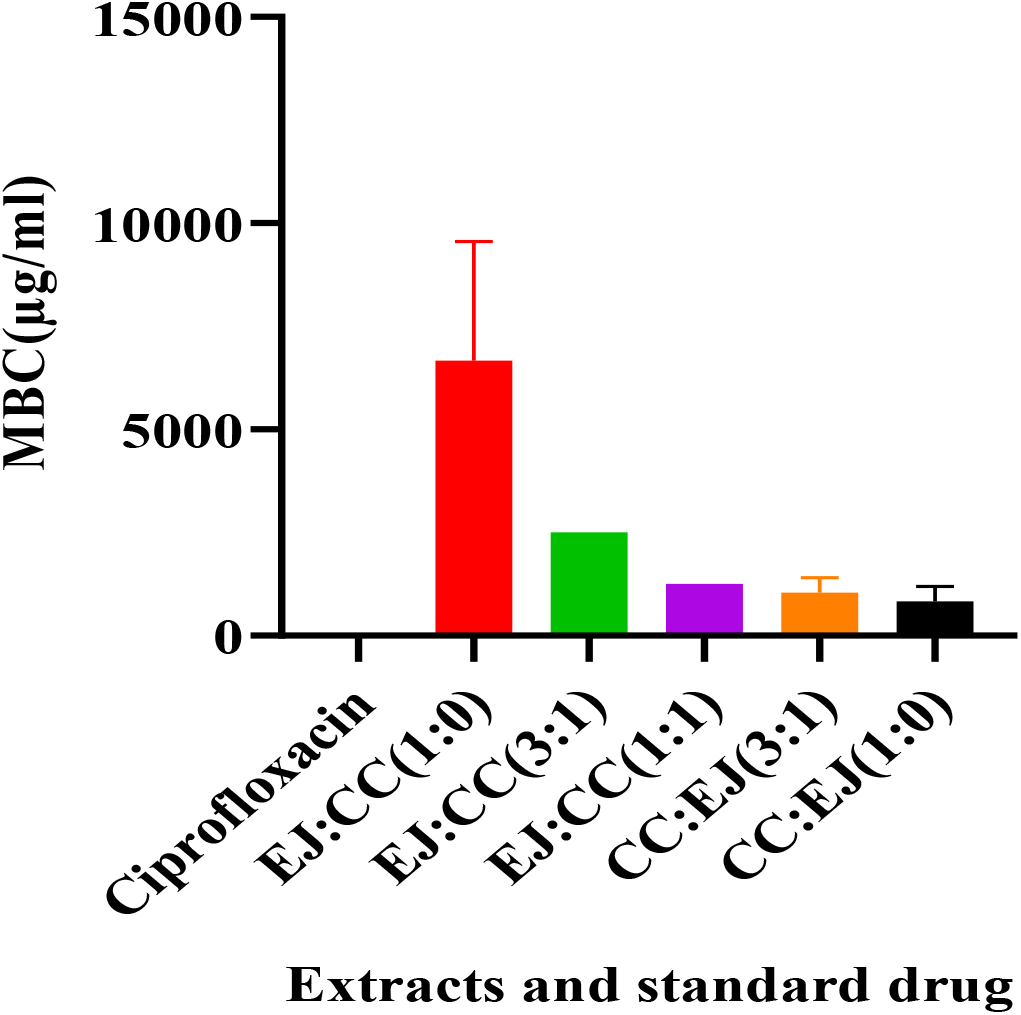
showing the MBCs of different extract proportions and standard drug.

### Antimicrobial interactions

The FICI/FBCI of the different extract proportions were between 1.5 and 3.917

## Discussion

The percentage yield of *Callistemon citrinus* and *Eriobotrya japonica* \were higher than the ones obtained by Krishna [21] for callistemon citrinus and [18] for *Eriobotrya japonica*. This could be attributed to the different methods of extraction, filtration and the difference in the percentage of ethanol. Krishna used Soxhlet Extraction whereas in our cold maceration was used in this particular study. Seong on the other hand used 5 % ethanol with Soxhlet extraction whereas in our study we used 60 % ethanol. Results of phytochemical screening were similar to Pendyala and Thaakur [28] for *Callistemon citrinus* and Hasibuan [29] for *Eriobotrya japonica* who also used ethanol for extraction.

The current results showed that the individual plant proportions possessed antimicrobial activity, with *Callistemon citrinus* being more potent than *Eriobotrya japonica.* This could be due to high quantity of Flavonoids in *Callistemon citrinus*. CC and EJ showed MIC and MBC values ranging from 521 to 3333 μg/ml and 1042 to 3667 μg/ml respectively. However, CC: EJ (1:0) had the lowest MIC and MBC comparable to that of the standard drug at P<0.05.

When determining the interactions according to Pratoomsoot [26], the FICI/FBCI was interpretated as follows; FICI/FBCI ≤0.5: synergy; 0.5 < FICI/FBCI ≤1: additivity; 1 < FICI/FBCI ≤4: no interaction (indifference); FICI/FBCI >4: antagonism. In the present study, the FICI/FBCI values obtained were 1 < FICI/FBCI ≤ 4 which revealed that there were no interactions between the two plant extracts.

## Conclusion

Therefore, CC: EJ (1:0) proportion is the best for formulating a standard alternative herbal product for prevention of dental carie*s*.

## Acknowledgment

We extend our most sincere appreciation to the Director Epicentre Mbarara, Dr. Juliet Mwanga Amumpaire for accepting us to carry out our research from their Microbiology laboratory. Special thanks also go to the Laboratory Manager Mr. Dan Nyehangane, Laboratory Technician Mr. Derrick Hope together with their team for the assistance rendered while carrying out our research at Epicentre. We also appreciate everyone at Mbarara University Pharmaceutical Chemistry Analytical Research Laboratory (MUPCARL) for their technical support.

We are also grateful to our supervisor Mr. Angupale Jimmy Ronald who guided us during the research period. Tremendous gratitude goes to every group member for exhibiting maximum cooperation.

